# A role of cellular translation regulation associated with toxic Huntingtin protein

**DOI:** 10.1101/687764

**Authors:** Hiranmay Joag, Vighnesh Ghatpande, Maitheli Sarkar, Anshu Raina, Meghal Desai, Mrunalini Shinde, Tania Bose, Amitabha Majumdar

## Abstract

Huntington’s disease (HD) is a severe neurodegenerative disorder caused by poly Q repeat expansion in the Huntingtin (Htt) gene. While the Htt amyloid aggregates are known to affect many cellular processes, its role in translation is not addressed. Here we report pathogenic Htt expression causes protein synthesis deficit in cells. We find a functional prion-like protein, the translation regulator Orb2 to be sequestered by Htt aggregates. Coexpression of Orb2 can partially rescue the lethality associated with poly Q expanded Htt. These findings can be relevant for HD as human homologs of Orb2 also can be sequestered by pathogenic Htt aggregates. Our work suggests that translation dysfunction could be one of the contributors in the pathogenesis of HD and new therapies targeting protein synthesis pathways might help alleviate disease symptoms.

## Introduction

Huntington’s disease (HD) is an autosomal dominant neurodegenerative disorder with no known treatment till date [3]. The symptoms associated with the disease are severe motor defects like chorea, uncoordinated movements, cognitive and behavioral problems which are progressive in nature. In the human brain, HD phenotypes are manifested as severe atrophy of caudate nucleus and putamen regions which together are part of the dorsal striatum. While the adult-onset HD phenotypes start manifesting after 40 years of age, there are also cases of juvenile HD which start manifesting at very early ages and are more aggressive in nature [47]. HD is caused due to CAG repeat expansion in the Exon1 of Huntingtin gene on chromosome 4p16 [39]. The CAG triplet codes for Glutamine (Q) and disease-causing expansions can range from 40 to 200 [36]. Due to polyQ expansion, the expanded Huntingtin (Htt) protein form misfolded aggregates causing an imbalance in cellular proteostasis and eventually neuronal cell death. Cellular proteostasis is regulated through a careful orchestration of protein synthesis, it’s folding and degradation [33]. In HD models, while the involvement of protein folding pathways through chaperone networks and proteasomal degradation pathways are well studied [5, 18, 20, 22, 35, 41, 51], its connection to protein synthesis is not very well established. There are several indicators pointing towards such a plausible connection. A network of proteins involved in rRNA processing and ribosome biogenesis were found to be differentially expressed in Yeast expressing pathogenic HttQ103 [57]. Components of ribosome quality control are also involved in Htt aggregate clearance in Yeast [60]. In mouse brains, both non-pathogenic and pathogenic Htt was found to be associated with translating polysomes [10]. A proteomics study with Neuro2a cells expressing pathogenic Htt, observed proteins involved in RNA processing, ribosome biogenesis, and translation to interact with oligomers of Htt [32]. Further proteomic profiling study of the progression of HD in mouse model also found several RNA binding proteins involved in translational control, ribosome biogenesis is part of the insoluble proteome in the brain [23], suggesting pathogenic Htt might be pushing these proteins to become non-functional. HD brains also show sense and antisense repeat-associated non-ATG (RAN) translation proteins which are also responsible for toxicity [2].

Motivated with these observations here we investigated the status of cellular translation associated with non-pathogenic and pathogenic Htt protein expression. We find pathogenic Htt expression causes cellular translation deficit in both Drosophila and Yeast models. We further find two isoforms of Orb2, a translation regulator in Drosophila gets sequestered by Htt aggregates. The sequestration probably happens through adsorption like process. Coexpression of Orb2 isoforms can rescue the lethality caused due to pathogenic Htt expression in Drosophila. We further find homologs of Orb2 in humans the Cytoplasmic polyadenylation binding proteins (hCPEB1-4) also can be sequestered by Htt aggregates, suggesting our observations with Orb2 might be relevant in HD.

## Results

### Expression of pathogenic HttQ138 is associated with reduced translation in cells

We expressed RFP tagged constructs of non-pathogenic HttQ15 and pathogenic HttQ138 in S2 cells. The Htt fragment used here codes for Caspase-6 cleaved N terminal 588 amino acid protein, is important for the disease progression as mutating this to prevent cleavage by Caspase-6 blocks neurodegeneration [16]. The HttQ15 construct is expressed in a diffused cytoplasmic pattern whereas the pathogenic Q138 shows punctate aggregates in cells. To investigate for translation in these cells we performed polysome analysis. Polysomes are complexes of ribosomes with the mRNAs it is translating to proteins [58]. For this cells are incubated with a translation inhibitor Cycloheximide which stalls the translating ribosomes on the mRNAs. Lysates from these cells are then centrifuged on a 5-45% sucrose density gradient which separates complexes based on their sedimentation coefficients, and as there is RNA in these complexes, absorbance measurement at 254 nm can be used to visualize the separation of the 40S, 60S, 80S ribosomes, and polysome fractions. For quantitation purpose, we compared the area under the curve for polysome and 80s and represent the data as a ratio between the two. In comparison to HttQ15, we observed a ∼24% decrease in this ratio for HttQ138 expressing cells suggesting that there are fewer polysomes or reduced translation associated with the pathogenic construct (Fig 1A, B).

**Figure 1:**
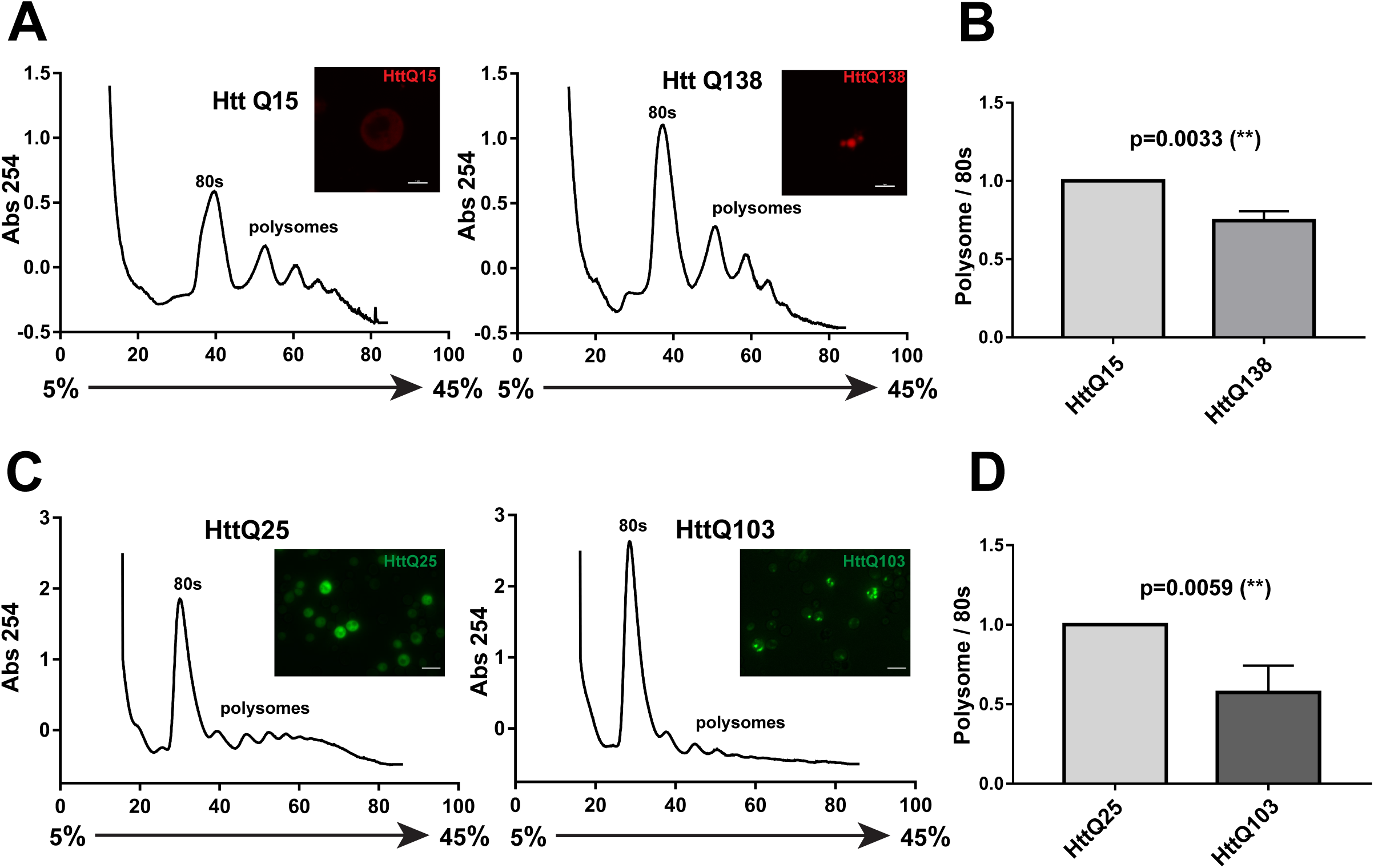
(A) Polysome profiles of S2 cells expressing HttQ15 and HttQ138. Inset show images of S2 cell expressing RFP tagged HttQ15 and HttQ138 **(B)** Quantitation of Polysome/80s ratio. Data is from n=4 experiments and is represented as relative fold change for HttQ138 compared to HttQ15. SEM is depicted as error bars. Significance is tested using unpaired one-tailed t-test and the p-value is 0.033(**) **(C)** Polysome profiles of yeast expressing HttQ25 and HttQ103. Inset show images of cells expressing GFP tagged HttQ25 and HttQ103 **(D)** Quantitation of Polysome/80s ratio from HttQ25 and HttQ103 expressing yeast. Data from n=3 experiments, represented as relative fold change of HttQ103 compared to HttQ25 and significance tested with unpaired one-tailed t-test and the p-value is 0.0059 (**). SEM is depicted as error bars.

We also performed polysome analysis In Yeast model of HD expressing Exon1 of Huntingtin with 25 Q (non-pathogenic) and 103Q repeats (pathogenic) under galactose-inducible promoter [43]. Similar to our observations with S2 cells, here we find ∼42.5% decreased polysome/80s ratio for pathogenic HttQ103 compared to non-pathogenic HttQ25 (Fig 1C, D). One common feature in the polysome profiles of both Drosophila S2 cells and Yeast is a higher 80s peak associated with pathogenic Htt expression. These results from two different systems suggest the expression of pathogenic Htt is associated with decreased polysome/80s ratio in cells.

We further used the Puromycin incorporation assay to compare levels of newly translated proteins in HttQ15 and HttQ138 cells. Puromycin gets incorporated in translating proteins and blocks further incorporation of amino acids and halts translation by premature chain termination [45]. The cells can then be lysed and puromycylated protein detected on western blots using anti Puromycin antibody [11, 54]. Here we observed ∼46.6% lesser Puromycin incorporation in HttQ138 cells in comparison to HttQ15 cells (Fig 2A, B). Together these observations suggest pathogenic Htt expression results in translation dysfunction in cells.

**Figure 2:**
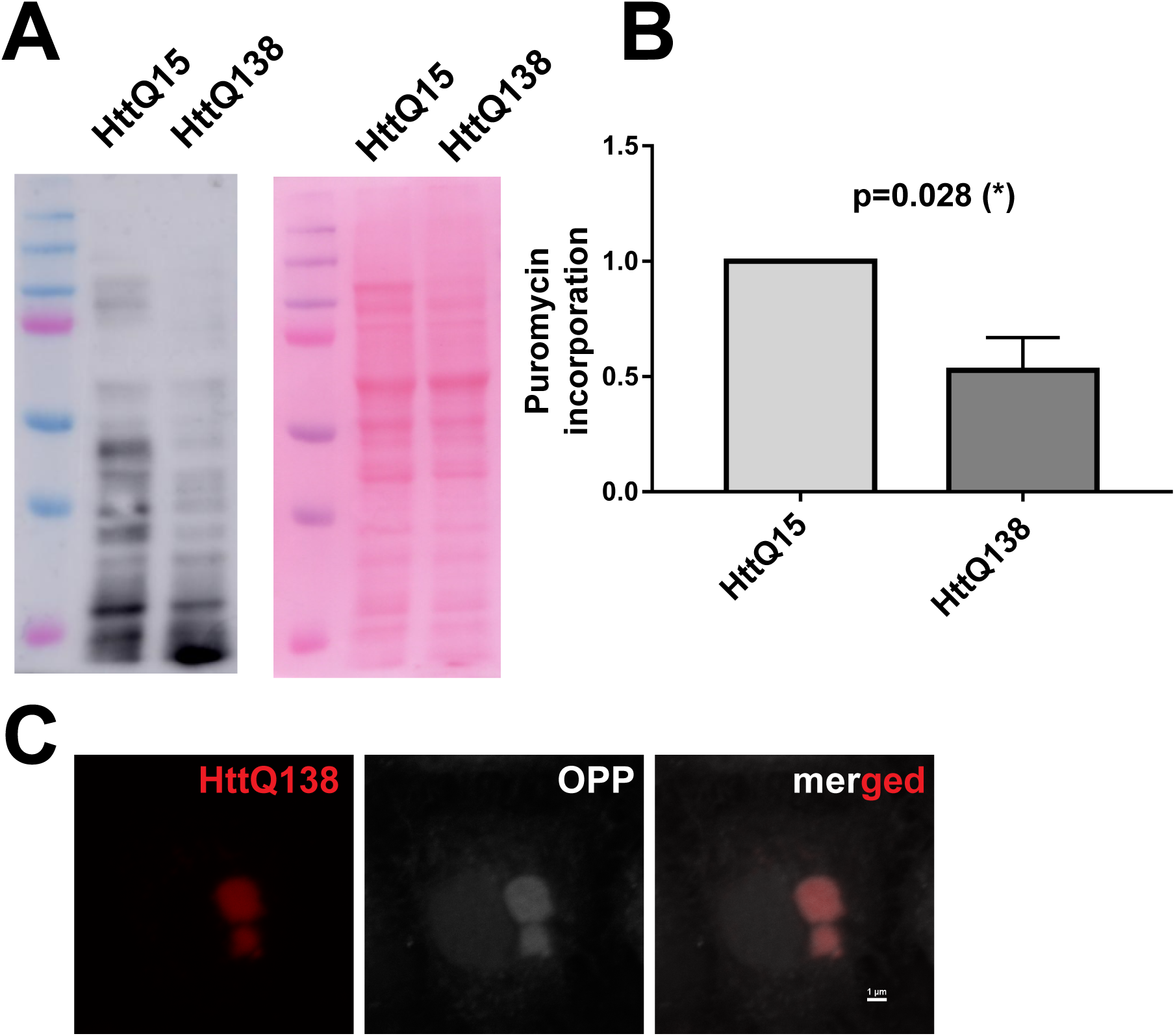
**(A)** Puromycin incorporation assay using anti Puromycin antibody, western blot shows reduced Puromycin incorporation in HttQ138 cells in comparison to HttQ15 cells. Right panel is Ponceau-S stained membrane for the same blot **(B)** Quantitation of Puromycin incorporation from HttQ15 and HttQ138 cells shows significantly reduced Puromycin incorporation in HttQ138 cells. Data is from n=3 experiments and is represented as relative fold change for HttQ138 compared to HttQ15. Significance is tested using an unpaired one-tailed t-test and the p-value is 0.028 (*). SEM is depicted as error bars (0.425±0.09). **(C)** O-Propargyl-Puromycin (OPP) incorporation assay shows incorporation of OPP in HttQ138 aggregates.

In order to visualize nascent protein synthesis in cells, we used an alkyne analog of Puromycin called O-propargyl-puromycin (OPP) which similar to Puromycin gets incorporated in translating proteins and can be detected by a click based chemical reaction [37]. Surprisingly, upon staining HttQ138 cells with a fluorescent OPP we noticed its incorporation in Htt aggregates (Fig 2C). Based on this observation and a previous study showing Htt binding to its own RNA [9] we hypothesized probably Htt aggregates are sequestering the RNA and translation machinery or its regulators and due to this some amount of translation, albeit stalled, can occur in the aggregates.

### Isoforms of a protein synthesis regulator Orb2 are sequestered by HttQ138 aggregates and rendered non-dynamic

As we noticed OPP incorporation in the Htt aggregates in the cell, we asked if other RNA binding proteins are also present in the aggregates. Previously Orb2A, an isoform of the translational regulator Orb2 was reported to be sequestered by HttQ128 in *Drosophila* larval brain [21]. The Htt construct used in this previous report was a shorter Htt fragment with lesser Q repeats. Orb2 a functional prion-like protein is the homolog of mammalian Cytoplasmic Polyadenylation Element Binding protein (CPEB) which can de-adenylate or polyadenylate the poly A tail of its target mRNAs by binding to the CPE elements in their 3’UTR [24]. Orb2 is necessary for the maintenance of long term memory [30, 40] and has two isoforms Orb2A and Orb2B, both containing an unstructured low complexity prion-like domain [40, 31]. On coexpressing GFP tagged Orb2A and Orb2B transgenes along with HttQ138 in the optic lobe neurons, we noticed both to colocalize with HttQ138 aggregates suggesting their sequestration by Htt aggregates (Fig 3A).

**Figure 3:**
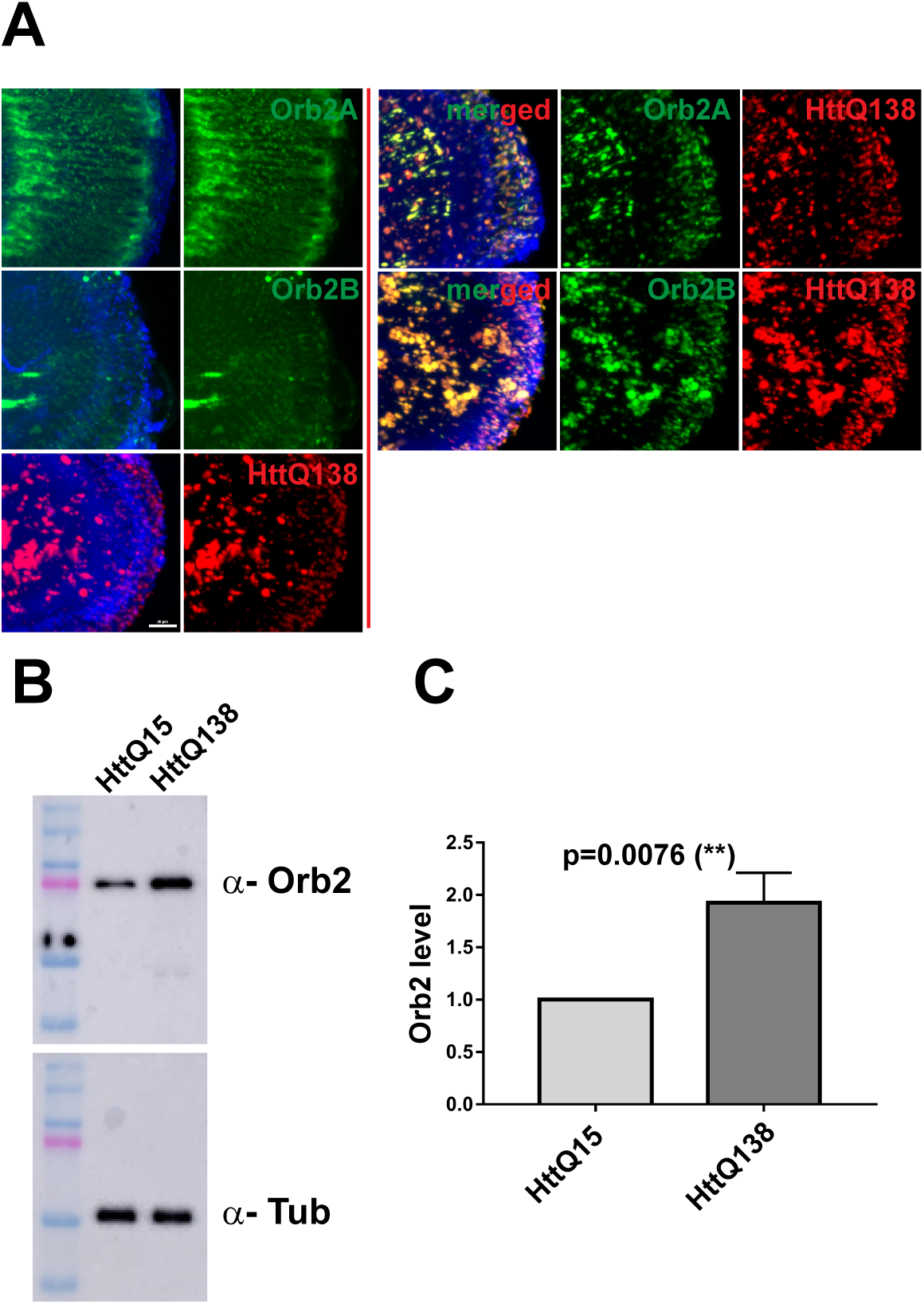
**(A)** Left panels show Drosophila optic lobes expressing GFP tagged Orb2A, Orb2B and RFP tagged HttQ138 with DAPI in Blue color. Right panels show optic lobes coexpressing Orb2A and Orb2B with HttQ138. In the case of coexpression, Orb2A and Orb2B are sequestered in the HttQ138 aggregates **(B)** Western blot from Drosophila head extracts of HttQ15 and HttQ138 shows increased levels of Orb2 associated with HttQ138. Anti-Tubulin antibody is used as the loading control. Quantitation of relative Orb2 levels shows increased Orb2 in HttQ138 expressing Drosophila heads. Data is from n=4 experiments and is represented as relative fold change for HttQ138 as compared to HttQ15. SEM is depicted as error bars. Significance is tested using an unpaired one-tailed t-test and the p-value is 0.0076 (**)

We next asked what happens to endogenous Orb2 levels in Htt Q15 and Q138 expressing fly heads. Of the two isoforms, Orb2B is readily detectable in the fly head extract due to its higher enrichment in comparison to Orb2A [40]. In western blots, we noticed higher amounts of Orb2B in HttQ138 fly heads compared to HttQ15 (Fig 3B, C). The increased levels might represent a steady state situation arising from compromised proteostasis situation in the brain or change in their half-life due to sequestration by Htt aggregates.

How is Orb2 different in its sequestered state in comparison to non-sequestered state? To address this we performed fluorescence recovery after photobleaching (FRAP) experiments with S2 cells coexpressing Orb2 with HttQ138 aggregates. In S2 cells also we observed the sequestration of both the Orb2 isoforms by Htt aggregates (Fig 4A). Compared to unsequestered Orb2A and Orb2B where a decreased but partial recovery for Orb2A and almost complete recovery for Orb2B could be seen, in the sequestered state with HttQ138 there was almost no recovery for both (Fig 4B, C). This indicates that the dynamic nature and probably function of Orb2 are lost in the sequestered state.

**Figure 4:**
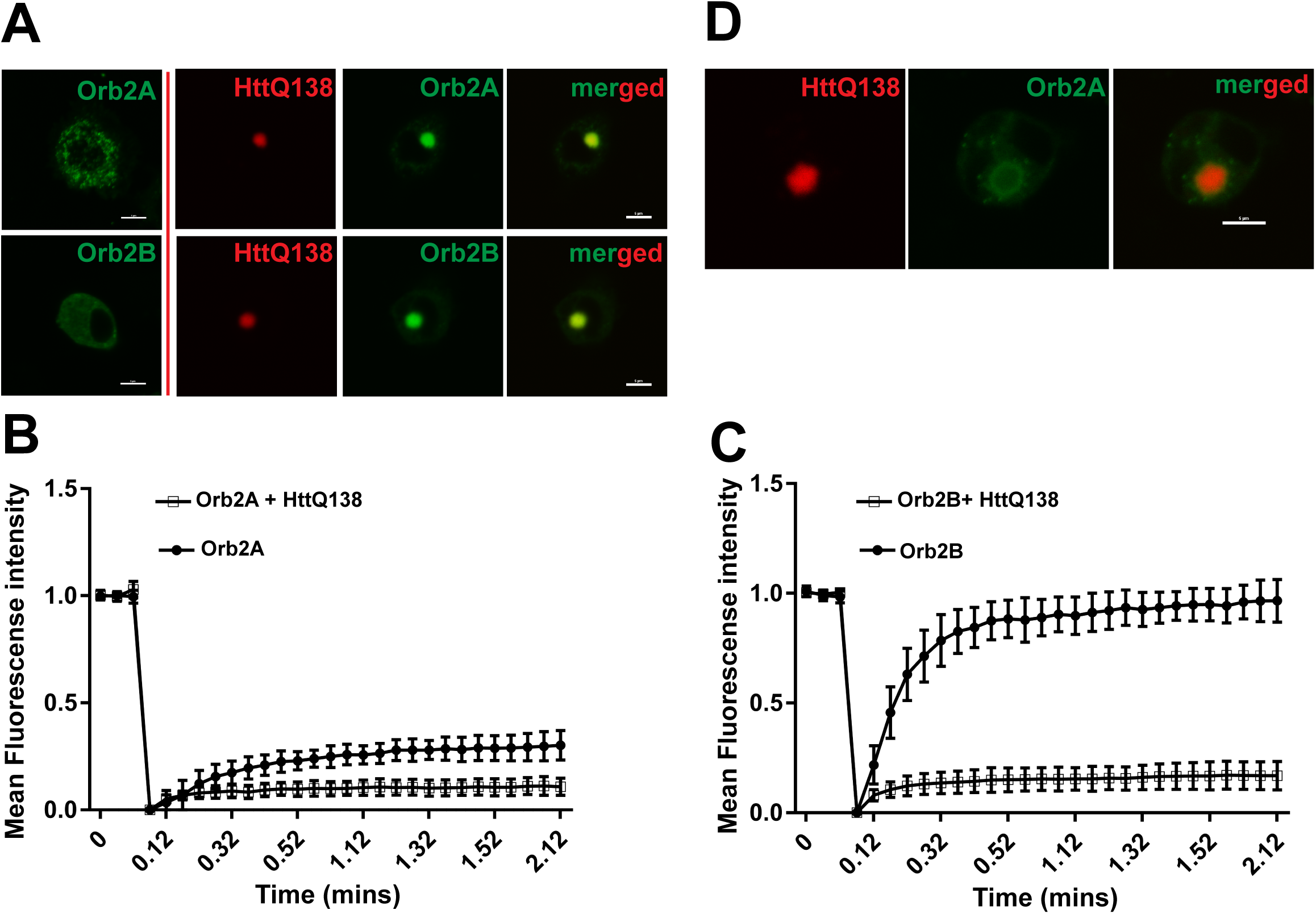
**(A)** S2 cells expressing Orb2A and Orb2B in left panels showing their expression all over the cell. Panels on right side of the red line shows sequestration of Orb2A and Orb2B on their coexpression with HttQ138 **(B & C)** Fluorescence recovery after photobleaching (FRAP) experiment data shows significantly decreased recovery for sequestered Orb2A and Orb2B in presence of HttQ138 in comparison to only Orb2A and Orb2B expressing cells. Data from n=10 experiments were tested using Mann-Whitney unpaired t-test and the p values are <0.0001 **(D)** Representative image of a fused cell fusion suggests sequestration of Orb2 in HttQ138 aggregates is through adsorption like process.

We further asked if the sequestration is due to coexpression of two sticky proteins inside the cell or through a seeding like mechanism. Towards this, we fused Orb2AGFP expressing cells with HttQ138 cells. In the fused cells, we observed the presence of Orb2 on the peripheral surface of Htt aggregates and not in their core (Fig 4D). These are probably representatives of starting intermediates in the process of sequestration of Orb2 by Htt aggregates using adsorption like process.

### Coexpression of Orb2 isoforms rescue lethality associated with pathogenic HttQ138

We wanted to know the effect of Orb2 coexpression on the phenotypes associated with Huntington’s disease model in Drosophila. Pan-neuronal expression of HttQ138 using Elav Gal4 causes lethality at pupal stages where only 4.8% of flies emerge. On coexpression of Orb2A and Orb2B, we found a rescue to the extent of 60 and 51% in the survival rate (Fig 5A). We next performed a loss of function study using an Orb2 RNAi line to test if decreasing Orb2 levels can increase HttQ138 associated lethality. Towards this end, we used a weaker expressing Elav Gal4 line to co-express HttQ15 and HttQ138 along with control Luciferase and Orb2 RNAi line. Low-level expression of HttQ15 with Orb2 RNAi and control Luciferase RNAi had no effect in the survival of animals. However on coexpression with HttQ138 we observed Orb2 RNAi reduced survival to 50% compared to around 77.8 % survival with Luciferase RNAi (Fig 5B). This suggests Orb2 knockdown increases the lethality associated with Htt aggregates. Together the gain and loss of function experiments suggest Orb2 genetically interacts with the Htt associated pathways and can modulate its toxicity.

**Figure 5:**
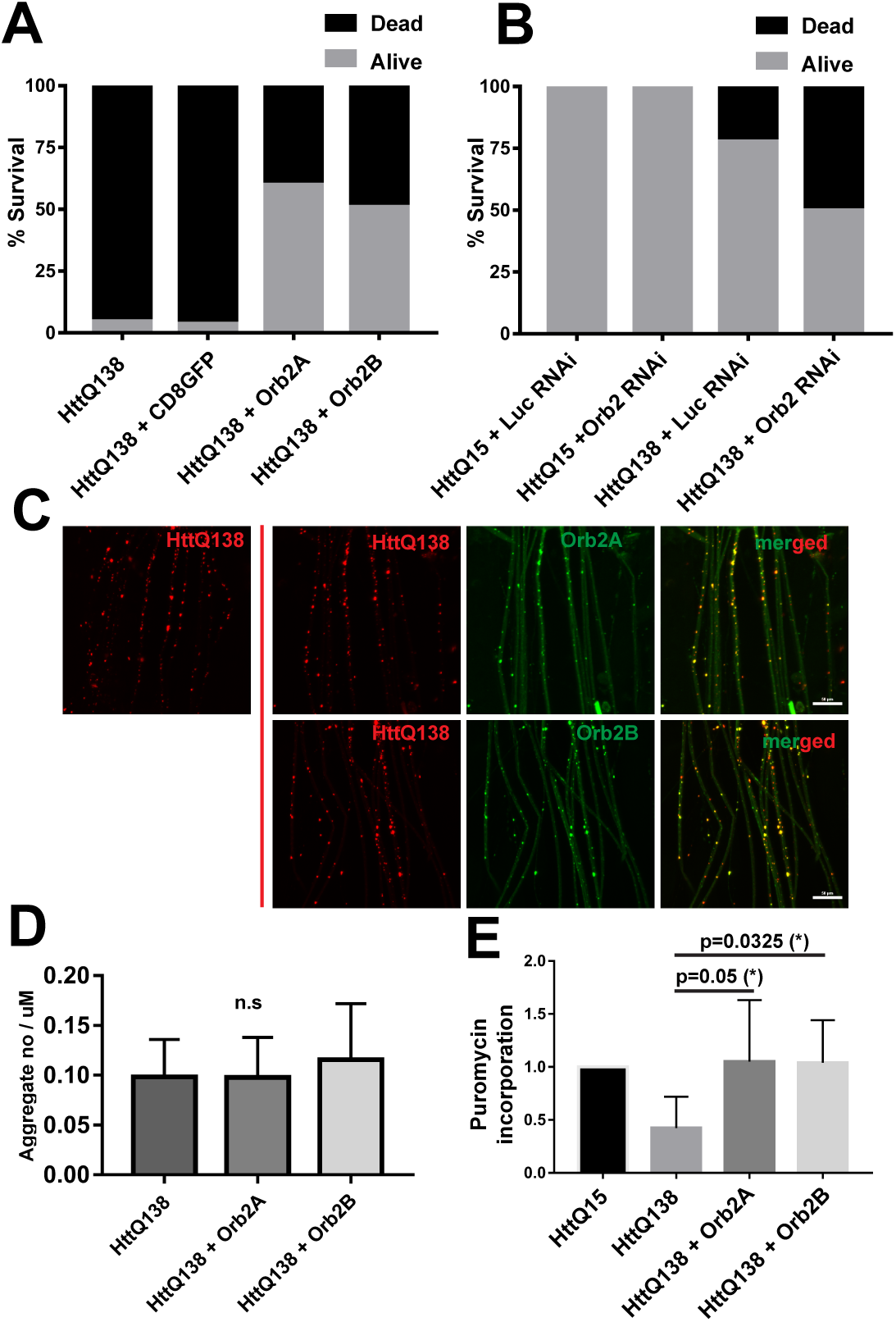
**(A)** Eclosion experiments show that coexpression of Orb2A and Orb2B can rescue the lethality associated with pan-neuronal expression of HttQ138 using Elav Gal4. In comparison to only HttQ138 (n=608) which showed 4.8% eclosion, coexpression with CD8GFP (n=26) showed 3.8 % eclosion and with Orb2A (n=131) and Orb2B (n=187) showed 60% and 51% eclosion respectively **(B)** Eclosion experiments performed with a lower expressing Elav Gal4 line show that in presence of Orb2RNAi coexpression there is decreased survival of HttQ138 expressing animals. For nonpathogenic HttQ15 coexpression with Luciferase (Luc) RNAi (n=35) and Orb2 RNAi (n=35) did not show any difference. HttQ138 coexpression with LucRNAi showed 77.8% survival (n=185) and with Orb2 RNAi showed 50% survival (n=165) **(C)** HttQ138 aggregate numbers in motor neurons in larvae expressing HttQ138 alone (left panel) and together with Orb2A and Orb2B (right panels) do not show any significant difference **(D)** Quantitation of aggregate numbers from HttQ138 alone (n=3, 11 axons) and HttQ138 with Orb2A (n=3, 20 axons) and Orb2B from (n=3, 18 axons) motoneurons do not show any significant difference when tested with Mann-Whitney unpaired t-test. (**E)** Quantitation of Puromycin incorporation from HttQ138 and HttQ138+ Orb2A GFP and HttQ138+ Orb2B GFP expressing cells shows significantly reduced Puromycin incorporation in HttQ138 cells. Data is from n=3 experiments and is represented as relative fold change for HttQ138 compared to HttQ15. Significance is tested using an unpaired one-tailed t-test and the p-value is 0.028 (*).

### Coexpression of Orb2 isoforms does not decrease the aggregate load but rescues the translation deficit

As coexpression of Orb2A and Orb2B could rescue the lethality associated with pathogenic HttQ138 we next asked what the mechanism of this rescue might be. As pathogenic Htt forms misfolded toxic aggregates, we first asked if coexpression of Orb2 isoforms is decreasing the Htt aggregate load in neurons. To test this we imaged the axons coming out of the ventral ganglion in Drosophila larvae. In animals coexpressing Orb2A and Orb2B with pathogenic HttQ138, we observed sequestration of these proteins in the Q138 aggregates (Fig 5C). This phenotype in the larval axons is similar to what we have observed in the optic lobe neurons and S2 cells. On quantitation of the Htt aggregates in axons however we found there is no significant difference between HttQ138 and the Orb2A, Orb2B rescued animals (Fig 5D), suggesting the mode of rescue by Orb2 coexpression is most likely not due to a decrease in aggregate load.

We next revisited our previous observation that translation is perturbed in HttQ138 cells and asked what happens on coexpression of Orb2 isoforms. We performed Puromycin incorporation experiment to quantitate protein synthesis and observed co-expression of Orb2A and Orb2B rescues the reduced translation associated with HttQ138 (Fig 5E). Overall these experiments suggest that increasing the translatory status of the cell is might mediate the rescue of the pathogenicity associated with HttQ138.

### Human CPEB’s can also be sequestered by Htt aggregates

Mammals including humans have 4 CPEB genes, CPEB1-4. We asked if human CPEB’s can also be sequestered by pathogenic Htt. We coexpressed GFP tagged hCPEB1-4 with HttQ138. In all cases, we observed the hCPEB’s to be sequestered by Htt aggregates (Fig 6A). On performing FRAP experiments we noted that similar to Orb2, hCPEB2-4 are rendered non-dynamic by sequestration in Htt aggregates. While there was some recovery of sequestered hCPEB1, but this is still significantly lower in comparison to only hCPEB1 (Fig6B). This suggests the hCPEB’s like their Drosophila homolog can be sequestered by Htt aggregates are probably made non-functional.

**Figure 6:**
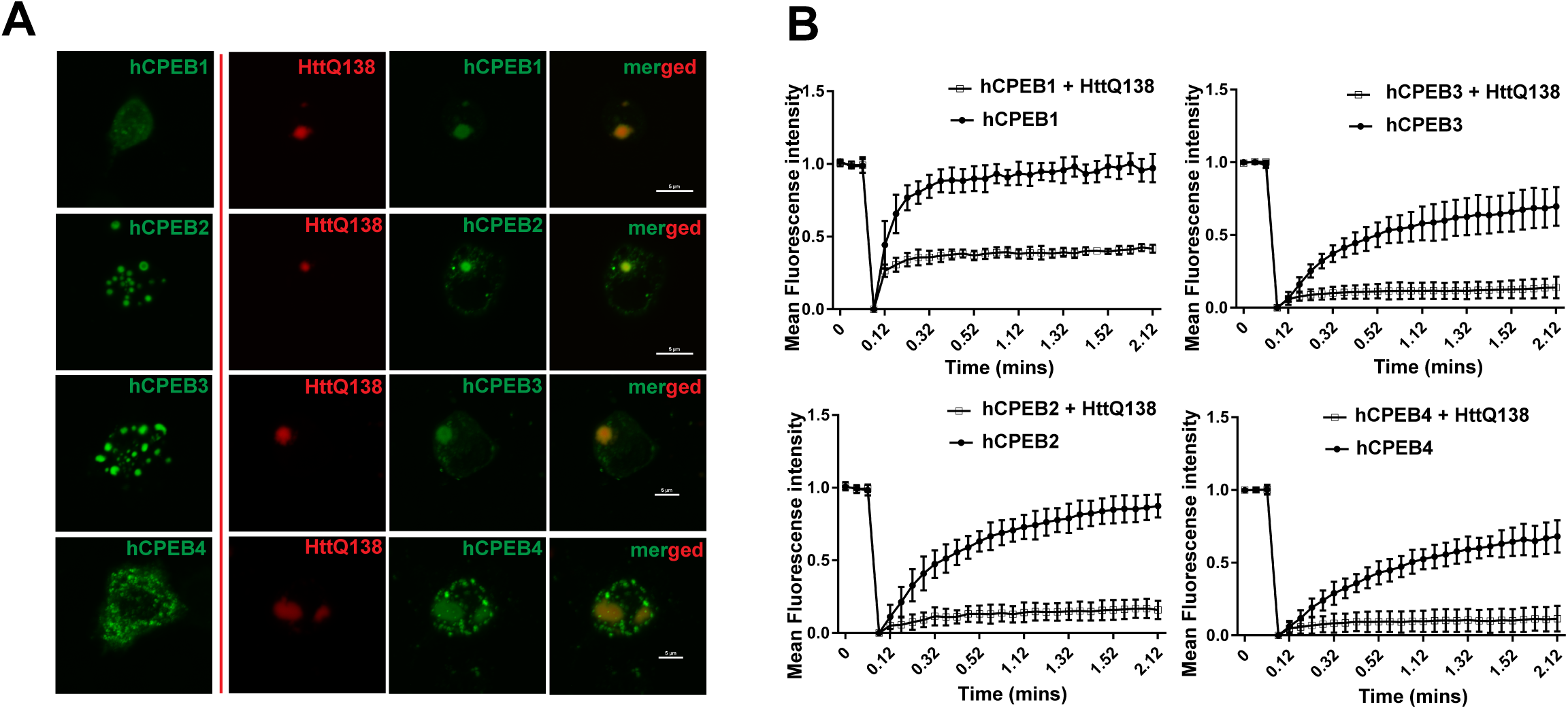
**(A)** S2 cells expressing Human CPEB’s (hCPEB1-4) in the left panel. Right panels show sequestration of Human CPEB’s on coexpression with HttQ138 **(B)** Fluorescence recovery after photobleaching (FRAP) experiment data shows significantly decreased recovery for sequestered hCPEB1, 2, 3 and 4 in the presence of HttQ138 in comparison to only hCPEB1-4 expressing cells. Data from n=9 experiments were tested using Mann-Whitney unpaired t-test and the p values are <0.0001.

## Discussions

Our observations suggest that pathogenic Htt expression can decrease translation in cells. Previous studies with Alzheimer’s, pathogenic Tau, prion disease and ALS/FTD spectrum disorders associated with Fus, Tdp-43 and C9orf72 expansion have been reported to show translation dysfunction [12, 19, 25, 26, 38, 42, 44, 53, 61]. Our study indicates pathogenic Huntingtin protein can also cause similar dysfunction. One reason for the translation dysfunction is possibly through sequestration of translation regulating proteins. Here we find that a translation regulator Orb2 can be one of the candidates which can be sequestered by Htt aggregates and thus can make cells deficient for Orb2 function. The sequestration of Orb2, a prion-like protein, by Htt aggregate resembles the cross-seeding phenomenon seen with other proteins associated with neurodegenerative conditions [14, 15, 17, 27, 46, 56]. Orb2 was previously shown to be a modifier of GGGGCC expansion disease model associated with C9orf72 and Ataxin-3 mediated neurodegeneration in Drosophila [6, 7]. Here we find coexpression of Orb2 rescues the Htt associated toxicity. What might be the reason for the rescue of HttQ138 associated toxicity by Orb2? One possibility is overexpression and sequestration of it by HttQ138 prevents the HttQ138 aggregates from sequestering other proteins. In the yeast model for HD, prion-like proteins were observed to rescue the toxicity associated with HttQ103 expression by reducing the sequestration of other prion-like proteins without decreasing the aggregate numbers [28, 52]. Here we notice no significant difference between the Htt aggregates in the axons of only HttQ138 expressing larvae and the Orb2A and B rescued ones. Another possibility is, as several ribosomal proteins and stress responsive kinases, chaperones, proteins involved in unfolded protein response have putative Orb2 binding elements in its 3’ UTR [55], Orb2 isoform overexpression can somehow modulate their expression to rescue the effects of their downregulation. Overexpression of two chaperones having CPE elements in there 3’ UTR, DnaJ1 and Mrj are previously reported to rescue Htt associated toxicity in Drosophila [13, 29].

Our findings connecting Orb2 with HD in the Drosophila model might also be relevant in humans as we find Htt aggregates can sequester hCPEB1-4. In the human genome, almost 20% of the genes can be CPEB1 targets [4, 49]. In mice model for HD, CPEB3 and 4 were present in the insoluble proteome in relevant brain regions [23] and CPEB4 specific target mRNAs were found to be enriched in deadenylated transcripts [48]. Patients suffering from HD are also reported to have problems with memory [1, 8]. As Orb2 was identified as a regulator of maintenance of memory, it’s human homologs might also have similar roles. Translation dysfunction associated with sequestration of CPEB’s might be one of the factors behind memory associated symptoms in HD patients. Orb2 is probably not the only candidate for Htt sequestration, and towards this, an exhaustive screen for all other RNA binding proteins might be needed in the future to identify other possible targets. The translation regulation pathways can be important targets towards improving the disease-associated symptoms.

## Materials and methods

### Drosophila strains and plasmids

UAS-RFP HttQ15, UAS-RFP HttQ138 transgenic lines, and plasmids were kind gifts from Prof. Troy Littleton’s lab [59]. Elav gal4 (X and 2nd chromosome insertions), UAS-Orb2A-GFP and UAS-Orb2B-GFP lines, Orb2 RNAi lines were kind gifts from Dr. Kausik Si’s lab. Yeast expressing plasmids for HttQ15 and HttQ103 were kind gifts from Prof. Michael Sherman’s lab.

### Cell culture

S2 cells were grown in Schneider’s media supplemented with 10% FBS. Transfections of plasmids were done with Effectene using the manufacturer’s protocol.

### Antibodies

Anti-Puromycin antibody was obtained from Kerafast. Secondary antibodies were obtained from CST. Drosophila Orb2 antibody was developed in our lab. Western blots were processed using chemiluminescence detected in GE AI 600 Imager.

### Polysome analysis

Polysome analysis was done using the Biocomp gradient station. 5-45% sucrose density gradient in resolving buffer consisting of 140 mM NaCl, 25mM Tris-Cl (pH:8), 10mM MgCl2) was poured using Biocomp gradient station. Each gradient was 11 ml in volume. S2 cells expressing the mentioned constructs were incubated with cycloheximide solution (final concentration of 20 ug/ml) for 20 minutes, before spinning them down at 3500 g for 10 minutes. The supernatant was removed and cells were immediately lysed with lysis buffer consisting of 300mM NaCl, 50mM Tris.Cl (pH:8), 10mM MgCl_2_, 1mM EGTA, 1% Triton-X100, 0.02% sodium deoxycholate. RNase inhibitor (Rnasin, Promega) was added to the lysate to a final concentration of 1 unit/100 ul. The lysate was spun at 10000 g at 4^°^C for 10 mins, and the supernatant was removed to a separate Eppendorf tube, RNA concentration across samples were normalized and equal quantities of RNA were loaded on each gradient. The gradients were loaded on a swing bucket SW41 rotor and were spun at 4°C at 27000 g speed for 3 hours. The gradients were then unloaded from the ultracentrifuge and were fractionated with simultaneous monitoring of OD at 254 nm using a Biocomp fractionator station. For quantitation purpose, the ratio of area under the curve of polysome/80s was measured.

### Puromycin Incorporation Assay

S2 cells transfected with RFP-Htt Q15, RFP-Htt Q138 and Orb2A-GFP or Orb2B-GFP as noted in the figures. 48hrs post transfection cells were treated with 100ug/ml Puromycin (Sigma) for 30mins and sorted using BD FACSAria™ III Standard Sorter. Sorted cells were then lysed in S2 lysis buffer (150mM NaCl, 10mM Tris pH 7.5 and 0.1% NP40) spun at 10000g for 10 mins. The protein concentration of the lysates was measured using the Bradford Assay and an equal amount of protein was used to perform western blot analysis. Membranes were probed with anti-Puromycin antibody Kerafast (EQ0001) and goat anti-mouse secondary antibody CST.

### OPP staining

OPP staining was done using Click-iT plus OPP Alexa-647 protein synthesis kit as per manufacturer’s protocol. Cells were incubated at 25oC for 30 minutes after adding 1:400 dilution of the Click-iT OPP reagent in Schneiders media. These cells were further washed 3 times with PBS and then fixed with 4% formaldehyde solution and then permeabilized with 0.5% Triton X-100 in PBS for 15 min. For the Click-iT reaction, cells were incubated in the dark at room temperature in the Click-iT reaction cocktail. After 30 min, samples were washed with Click-iT reaction rinse buffer and PBS, followed by imaging.

### Cell fusion assay

Cell fusion assays were done by mixing cells expressing RFP tagged HttQ138 with Orb2A GFP expressing cells. These cells were coexpressing *C*. *elegans* fusogenic protein EFF1. Post mixing, fused cells were identified manually under a microscope and imaged within 4 hours.

### FRAP assay

FRAP experiments were performed using a 100X oil immersion lens in a Nikon Eclipse Ti based confocal system. The analysis was done using Fiji and easyfrap (https://easyfrap.vmnet.upatras.gr/) [34, 50].

## Author contributions

H.J started this project and identified Orb2 isoforms as rescuers of Htt pathogenicity. V.G performed the S2 cell polysome and puromycin incorporation experiments. M.S performed S2 cell imaging and FRAP experiments. M.S did the yeast polysome experiments. A.R did the Orb2 rescue and Orb2 RNAi experiments. M.D performed the Orb2 levels quantitations and OPP experiments. T.B and A.M conceived and designed the project. AM wrote the manuscript.

## Acknowledgments

We acknowledge Prof. Troy Littleton, Kausik Si for sharing several Drosophila stocks and plasmids with us. We also thank Addgene and Bloomington Drosophila stock center for some plasmids and lines used in this work. We thank Kausik Si, Gunther Hollopeter, and Deepa Subramanyam for their comments on the work. This work was supported by funding from Wellcome Trust-DBT India Alliance intermediate fellowship (IA/I/13/2/501030) to AM along with intramural funding from NCCS. TB was supported by Ramalingaswamy fellowship from DBT (BT/RLF/Re-entry/54/2013) and IYBA grant (BT/09/IYBA/2015/03).

## References

1. Aldaz T, Nigro P, Sánchez-Gómez A, Painous C, Planellas L, Santacruz P, Cámara A, Compta Y, Valldeoriola F, Martí MJ, Muñoz E (2019) Non-motor symptoms in Huntington’s disease: a comparative study with Parkinson’s disease. J Neurol 266:1340–1350. doi: 10.1007/s00415-019-09263-7

2. Bañez-Coronel M, Ayhan F, Tarabochia AD, Zu T, Perez BA, Tusi SK, Pletnikova O, Borchelt DR, Ross CA, Margolis RL, Yachnis AT, Troncoso JC, Ranum LPW (2015) RAN Translation in Huntington Disease. Neuron 88:667–677. doi: 10.1016/j.neuron.2015.10.038

3. Bates GP, Dorsey R, Gusella JF, Hayden MR, Kay C, Leavitt BR, Nance M, Ross CA, Scahill RI, Wetzel R, Wild EJ, Tabrizi SJ (2015) Huntington disease. Nat Rev Dis Primers 1:15005. doi: 10.1038/nrdp.2015.5

4. Belloc E, Piqué M, Méndez R (2008) Sequential waves of polyadenylation and deadenylation define a translation circuit that drives meiotic progression. Biochem Soc Trans 36:665–670. doi: 10.1042/BST0360665

5. Bennett EJ, Shaler TA, Woodman B, Ryu K-Y, Zaitseva TS, Becker CH, Bates GP, Schulman H, Kopito RR (2007) Global changes to the ubiquitin system in Huntington’s disease. Nature 448:704–708. doi: 10.1038/nature06022

6. Bilen J, Bonini NM (2007) Genome-Wide Screen for Modifiers of Ataxin-3 Neurodegeneration in Drosophila. PLOS Genetics 3:e177. doi: 10.1371/journal.pgen.0030177

7. Burguete AS, Almeida S, Gao F-B, Kalb R, Akins MR, Bonini NM (2015) GGGGCC microsatellite RNA is neuritically localized, induces branching defects, and perturbs transport granule function. eLife 4:e08881. doi: 10.7554/eLife.08881

8. Carmichael AM, Irish M, Glikmann-Johnston Y, Singh P, Stout JC (2019) Pervasive autobiographical memory impairments in Huntington’s disease. Neuropsychologia 127:123–130. doi: 10.1016/j.neuropsychologia.2019.02.017

9. Culver BP, DeClercq J, Dolgalev I, Yu MS, Ma B, Heguy A, Tanese N (2016) Huntington’s Disease Protein Huntingtin Associates with its own mRNA. J Huntingtons Dis 5:39–51. doi: 10.3233/JHD-150177

10. Culver BP, Savas JN, Park SK, Choi JH, Zheng S, Zeitlin SO, Yates JR, Tanese N (2012) Proteomic analysis of wild-type and mutant huntingtin-associated proteins in mouse brains identifies unique interactions and involvement in protein synthesis. J Biol Chem 287:21599–21614. doi: 10.1074/jbc.M112.359307

11. Deliu LP, Ghosh A, Grewal SS (2017) Investigation of protein synthesis in Drosophila larvae using puromycin labelling. Biol Open 6:1229–1234. doi: 10.1242/bio.026294

12. Ding Q, Markesbery WR, Chen Q, Li F, Keller JN (2005) Ribosome dysfunction is an early event in Alzheimer’s disease. J Neurosci 25:9171–9175. doi: 10.1523/JNEUROSCI.3040-05.2005

13. Fayazi Z, Ghosh S, Marion S, Bao X, Shero M, Kazemi-Esfarjani P (2006) A Drosophila ortholog of the human MRJ modulates polyglutamine toxicity and aggregation. Neurobiol Dis 24:226–244. doi: 10.1016/j.nbd.2006.06.015

14. Furukawa Y, Kaneko K, Matsumoto G, Kurosawa M, Nukina N (2009) Cross-Seeding Fibrillation of Q/N-Rich Proteins Offers New Pathomechanism of Polyglutamine Diseases. J Neurosci 29:5153–5162. doi: 10.1523/JNEUROSCI.0783-09.2009

15. Giasson BI, Forman MS, Higuchi M, Golbe LI, Graves CL, Kotzbauer PT, Trojanowski JQ, Lee VM-Y (2003) Initiation and synergistic fibrillization of tau and alpha-synuclein. Science 300:636–640. doi: 10.1126/science.1082324

16. Graham RK, Deng Y, Slow EJ, Haigh B, Bissada N, Lu G, Pearson J, Shehadeh J, Bertram L, Murphy Z, Warby SC, Doty CN, Roy S, Wellington CL, Leavitt BR, Raymond LA, Nicholson DW, Hayden MR (2006) Cleavage at the caspase-6 site is required for neuronal dysfunction and degeneration due to mutant huntingtin. Cell 125:1179–1191. doi: 10.1016/j.cell.2006.04.026

17. Guo JL, Covell DJ, Daniels JP, Iba M, Stieber A, Zhang B, Riddle DM, Kwong LK, Xu Y, Trojanowski JQ, Lee VMY (2013) Distinct α-synuclein strains differentially promote tau inclusions in neurons. Cell 154:103–117. doi: 10.1016/j.cell.2013.05.057

18. Hageman J, Rujano MA, van Waarde MAWH, Kakkar V, Dirks RP, Govorukhina N, Oosterveld-Hut HMJ, Lubsen NH, Kampinga HH (2010) A DNAJB chaperone subfamily with HDAC-dependent activities suppresses toxic protein aggregation. Mol Cell 37:355–369. doi: 10.1016/j.molcel.2010.01.001

19. Hartmann H, Hornburg D, Czuppa M, Bader J, Michaelsen M, Farny D, Arzberger T, Mann M, Meissner F, Edbauer D (2018) Proteomics and C9orf72 neuropathology identify ribosomes as poly-GR/PR interactors driving toxicity. Life Sci Alliance 1. doi: 10.26508/lsa.201800070

20. Hay DG, Sathasivam K, Tobaben S, Stahl B, Marber M, Mestril R, Mahal A, Smith DL, Woodman B, Bates GP (2004) Progressive decrease in chaperone protein levels in a mouse model of Huntington’s disease and induction of stress proteins as a therapeutic approach. Hum Mol Genet 13:1389–1405. doi: 10.1093/hmg/ddh144

21. Hervás R, Li L, Majumdar A, Fernández-Ramírez MDC, Unruh JR, Slaughter BD, Galera-Prat A, Santana E, Suzuki M, Nagai Y, Bruix M, Casas-Tintó S, Menéndez M, Laurents DV, Si K, Carrión-Vázquez M (2016) Molecular Basis of Orb2 Amyloidogenesis and Blockade of Memory Consolidation. PLoS Biol 14:e1002361. doi: 10.1371/journal.pbio.1002361

22. Hipp MS, Patel CN, Bersuker K, Riley BE, Kaiser SE, Shaler TA, Brandeis M, Kopito RR (2012) Indirect inhibition of 26S proteasome activity in a cellular model of Huntington’s disease. J Cell Biol 196:573–587. doi: 10.1083/jcb.201110093

23. Hosp F, Gutiérrez-Ángel S, Schaefer MH, Cox J, Meissner F, Hipp MS, Hartl F-U, Klein R, Dudanova I, Mann M (2017) Spatiotemporal Proteomic Profiling of Huntington’s Disease Inclusions Reveals Widespread Loss of Protein Function. Cell Rep 21:2291–2303. doi: 10.1016/j.celrep.2017.10.097

24. Ivshina M, Lasko P, Richter JD (2014) Cytoplasmic polyadenylation element binding proteins in development, health, and disease. Annu Rev Cell Dev Biol 30:393–415. doi: 10.1146/annurev-cellbio-101011-155831

25. Kamelgarn M, Chen J, Kuang L, Jin H, Kasarskis EJ, Zhu H (2018) ALS mutations of FUS suppress protein translation and disrupt the regulation of nonsense-mediated decay. Proc Natl Acad Sci USA 115:E11904–E11913. doi: 10.1073/pnas.1810413115

26. Kanekura K, Yagi T, Cammack AJ, Mahadevan J, Kuroda M, Harms MB, Miller TM, Urano F (2016) Poly-dipeptides encoded by the C9ORF72 repeats block global protein translation. Hum Mol Genet 25:1803–1813. doi: 10.1093/hmg/ddw052

27. Katorcha E, Makarava N, Lee YJ, Lindberg I, Monteiro MJ, Kovacs GG, Baskakov IV (2017) Cross-seeding of prions by aggregated α-synuclein leads to transmissible spongiform encephalopathy. PLoS Pathog 13:e1006563. doi: 10.1371/journal.ppat.1006563

28. Kayatekin C, Matlack KES, Hesse WR, Guan Y, Chakrabortee S, Russ J, Wanker EE, Shah JV, Lindquist S (2014) Prion-like proteins sequester and suppress the toxicity of huntingtin exon 1. Proc Natl Acad Sci USA 111:12085–12090. doi: 10.1073/pnas.1412504111

29. Kazemi-Esfarjani P, Benzer S (2000) Genetic suppression of polyglutamine toxicity in Drosophila. Science 287:1837–1840. doi: 10.1126/science.287.5459.1837

30. Keleman K, Krüttner S, Alenius M, Dickson BJ (2007) Function of the Drosophila CPEB protein Orb2 in long-term courtship memory. Nature Neuroscience 10:1587–1593. doi: 10.1038/nn1996

31. Khan MR, Li L, Pérez-Sánchez C, Saraf A, Florens L, Slaughter BD, Unruh JR, Si K (2015) Amyloidogenic Oligomerization Transforms Drosophila Orb2 from a Translation Repressor to an Activator. Cell 163:1468–1483. doi: 10.1016/j.cell.2015.11.020

32. Kim YE, Hosp F, Frottin F, Ge H, Mann M, Hayer-Hartl M, Hartl FU (2016) Soluble Oligomers of PolyQ-Expanded Huntingtin Target a Multiplicity of Key Cellular Factors. Mol Cell 63:951–964. doi: 10.1016/j.molcel.2016.07.022

33. Klaips CL, Jayaraj GG, Hartl FU (2018) Pathways of cellular proteostasis in aging and disease. J Cell Biol 217:51–63. doi: 10.1083/jcb.201709072

34. Koulouras G, Panagopoulos A, Rapsomaniki MA, Giakoumakis NN, Taraviras S, Lygerou Z (2018) EasyFRAP-web: a web-based tool for the analysis of fluorescence recovery after photobleaching data. Nucleic Acids Res 46:W467–W472. doi: 10.1093/nar/gky508

35. Labbadia J, Novoselov SS, Bett JS, Weiss A, Paganetti P, Bates GP, Cheetham ME (2012) Suppression of protein aggregation by chaperone modification of high molecular weight complexes. Brain 135:1180–1196. doi: 10.1093/brain/aws022

36. Lee J-M, Ramos EM, Lee J-H, Gillis T, Mysore JS, Hayden MR, Warby SC, Morrison P, Nance M, Ross CA, Margolis RL, Squitieri F, Orobello S, Di Donato S, Gomez-Tortosa E, Ayuso C, Suchowersky O, Trent RJA, McCusker E, Novelletto A, Frontali M, Jones R, Ashizawa T, Frank S, Saint-Hilaire MH, Hersch SM, Rosas HD, Lucente D, Harrison MB, Zanko A, Abramson RK, Marder K, Sequeiros J, Paulsen JS, PREDICT-HD study of the Huntington Study Group (HSG), Landwehrmeyer GB, REGISTRY study of the European Huntington’s Disease Network, Myers RH, HD-MAPS Study Group, MacDonald ME, Gusella JF, COHORT study of the HSG (2012) CAG 1 repeat expansion in Huntington disease determines age at onset in a fully dominant fashion. Neurology 78:690–695. doi: 10.1212/WNL.0b013e318249f683

37. Liu J, Xu Y, Stoleru D, Salic A (2012) Imaging protein synthesis in cells and tissues with an alkyne analog of puromycin. Proc Natl Acad Sci USA 109:413–418. doi: 10.1073/pnas.1111561108

38. López-Erauskin J, Tadokoro T, Baughn MW, Myers B, McAlonis-Downes M, Chillon-Marinas C, Asiaban JN, Artates J, Bui AT, Vetto AP, Lee SK, L. AV, Sun Y, Jambeau M, Boubaker J, Swing D, Qiu J, Hicks GG, Ouyang Z, Fu X-D, Tessarollo L, Ling S-C, Parone PA, Shaw CE, Marsala M, Lagier-Tourenne C, Cleveland DW, Da Cruz S (2018) ALS/FTD-Linked Mutation in FUS Suppresses Intra-axonal Protein Synthesis and Drives Disease Without Nuclear Loss-of-Function of FUS. Neuron 100:816-830.e7. doi: 10.1016/j.neuron.2018.09.044

39. MacDonald ME, Ambrose CM, Duyao MP, Myers RH, Lin C, Srinidhi L, Barnes G, Taylor SA, James M, Groot N, MacFarlane H, Jenkins B, Anderson MA, Wexler NS, Gusella JF, Bates GP, Baxendale S, Hummerich H, Kirby S, North M, Youngman S, Mott R, Zehetner G, Sedlacek Z, Poustka A, Frischauf A-M, Lehrach H, Buckler AJ, Church D, Doucette-Stamm L, O’Donovan MC, Riba-Ramirez L, Shah M, Stanton VP, Strobel SA, Draths KM, Wales JL, Dervan P, Housman DE, Altherr M, Shiang R, Thompson L, Fielder T, Wasmuth JJ, Tagle D, Valdes J, Elmer L, Allard M, Castilla L, Swaroop M, Blanchard K, Collins FS, Snell R, Holloway T, Gillespie K, Datson N, Shaw D, Harper PS (1993) A novel gene containing a trinucleotide repeat that is expanded and unstable on Huntington’s disease chromosomes. Cell 72:971–983. doi: 10.1016/0092-8674(93)90585-E

40. Majumdar A, Cesario WC, White-Grindley E, Jiang H, Ren F, Khan MR, Li L, Choi EM-L, Kannan K, Guo F, Unruh J, Slaughter B, Si K (2012) Critical role of amyloid-like oligomers of Drosophila Orb2 in the persistence of memory. Cell 148:515–529. doi: 10.1016/j.cell.2012.01.004

41. Martinez-Vicente M, Talloczy Z, Wong E, Tang G, Koga H, Kaushik S, de Vries R, Arias E, Harris S, Sulzer D, Cuervo AM (2010) Cargo recognition failure is responsible for inefficient autophagy in Huntington’s disease. Nat Neurosci 13:567–576. doi: 10.1038/nn.2528

42. Meier S, Bell M, Lyons DN, Rodriguez-Rivera J, Ingram A, Fontaine SN, Mechas E, Chen J, Wolozin B, LeVine H, Zhu H, Abisambra JF (2016) Pathological Tau Promotes Neuronal Damage by Impairing Ribosomal Function and Decreasing Protein Synthesis. J Neurosci 36:1001–1007. doi: 10.1523/JNEUROSCI.3029-15.2016

43. Meriin AB, Zhang X, He X, Newnam GP, Chernoff YO, Sherman MY (2002) Huntingtin toxicity in yeast model depends on polyglutamine aggregation mediated by a prion-like protein Rnq1. J Cell Biol 157:997–1004. doi: 10.1083/jcb.200112104

44. Moens TG, Niccoli T, Wilson KM, Atilano ML, Birsa N, Gittings LM, Holbling BV, Dyson MC, Thoeng A, Neeves J, Glaria I, Yu L, Bussmann J, Storkebaum E, Pardo M, Choudhary JS, Fratta P, Partridge L, Isaacs AM (2019) C9orf72 arginine-rich dipeptide proteins interact with ribosomal proteins in vivo to induce a toxic translational arrest that is rescued by eIF1A. Acta Neuropathol 137:487–500. doi: 10.1007/s00401-018-1946-4

45. Nathans D (1964) Puromycin Inhibition of Protein Synthesis: Incorporation of Puromycin into Peptide Chains. PNAS 51:585–592. doi: 10.1073/pnas.51.4.585

46. Ono K, Takahashi R, Ikeda T, Yamada M (2012) Cross-seeding effects of amyloid β-protein and α-synuclein. J Neurochem 122:883–890. doi: 10.1111/j.1471-4159.2012.07847.x

47. Orr HT, Zoghbi HY (2007) Trinucleotide repeat disorders. Annu Rev Neurosci 30:575–621. doi: 10.1146/annurev.neuro.29.051605.113042

48. Parras A, Anta H, Santos-Galindo M, Swarup V, Elorza A, Nieto-González JL, Picó S, Hernández IH, Díaz-Hernández JI, Belloc E, Rodolosse A, Parikshak NN, Peñagarikano O, Fernández-Chacón R, Irimia M, Navarro P, Geschwind DH, Méndez R, Lucas JJ (2018) Autism-like phenotype and risk gene mRNA deadenylation by CPEB4 mis-splicing. Nature 560:441–446. doi: 10.1038/s41586-018-0423-5

49. Piqué M, López JM, Foissac S, Guigó R, Méndez R (2008) A combinatorial code for CPE-mediated translational control. Cell 132:434–448. doi: 10.1016/j.cell.2007.12.038

50. Rapsomaniki MA, Kotsantis P, Symeonidou I-E, Giakoumakis N-N, Taraviras S, Lygerou Z (2012) easyFRAP: an interactive, easy-to-use tool for qualitative and quantitative analysis of FRAP data. Bioinformatics 28:1800–1801. doi: 10.1093/bioinformatics/bts241

51. Reis SD, Pinho BR, Oliveira JMA (2017) Modulation of Molecular Chaperones in Huntington’s Disease and Other Polyglutamine Disorders. Mol Neurobiol 54:5829–5854. doi: 10.1007/s12035-016-0120-z

52. Ripaud L, Chumakova V, Antonin M, Hastie AR, Pinkert S, Körner R, Ruff KM, Pappu RV, Hornburg D, Mann M, Hartl FU, Hipp MS (2014) Overexpression of Q-rich prion-like proteins suppresses polyQ cytotoxicity and alters the polyQ interactome. Proc Natl Acad Sci USA 111:18219–18224. doi: 10.1073/pnas.1421313111

53. Russo A, Scardigli R, La Regina F, Murray ME, Romano N, Dickson DW, Wolozin B, Cattaneo A, Ceci M (2017) Increased cytoplasmic TDP-43 reduces global protein synthesis by interacting with RACK1 on polyribosomes. Hum Mol Genet 26:1407–1418. doi: 10.1093/hmg/ddx035

54. Schmidt EK, Clavarino G, Ceppi M, Pierre P (2009) SUnSET, a nonradioactive method to monitor protein synthesis. Nat Methods 6:275–277. doi: 10.1038/nmeth.1314

55. Stepien BK, Oppitz C, Gerlach D, Dag U, Novatchkova M, Krüttner S, Stark A, Keleman K (2016) RNA-binding profiles of Drosophila CPEB proteins Orb and Orb2. PNAS 113:E7030–E7038. doi: 10.1073/pnas.1603715113

56. Tanaka M, Ishizuka K, Nekooki-Machida Y, Endo R, Takashima N, Sasaki H, Komi Y, Gathercole A, Huston E, Ishii K, Hui KK-W, Kurosawa M, Kim S-H, Nukina N, Takimoto E, Houslay MD, Sawa A (2017) Aggregation of scaffolding protein DISC1 dysregulates phosphodiesterase 4 in Huntington’s disease. J Clin Invest 127:1438–1450. doi: 10.1172/JCI85594

57. Tauber E, Miller-Fleming L, Mason RP, Kwan W, Clapp J, Butler NJ, Outeiro TF, Muchowski PJ, Giorgini F (2011) Functional gene expression profiling in yeast implicates translational dysfunction in mutant huntingtin toxicity. J Biol Chem 286:410–419. doi: 10.1074/jbc.M110.101527

58. Warner JR, Knopf PM, Rich A (1963) A multiple ribosomal structure in protein synthesis. Proc Natl Acad Sci USA 49:122–129. doi: 10.1073/pnas.49.1.122

59. Weiss KR, Kimura Y, Lee W-CM, Littleton JT (2012) Huntingtin aggregation kinetics and their pathological role in a Drosophila Huntington’s disease model. Genetics 190:581–600. doi: 10.1534/genetics.111.133710

60. Yang J, Hao X, Cao X, Liu B, Nyström T (2016) Spatial sequestration and detoxification of Huntingtin by the ribosome quality control complex. Elife 5. doi: 10.7554/eLife.11792

61. Zhang Y-J, Gendron TF, Ebbert MTW, O’Raw AD, Yue M, Jansen-West K, Zhang X, Prudencio M, Chew J, Cook CN, Daughrity LM, Tong J, Song Y, Pickles SR, Castanedes-Casey M, Kurti A, Rademakers R, Oskarsson B, Dickson DW, Hu W, Gitler AD, Fryer JD, Petrucelli L (2018) Poly(GR) impairs protein translation and stress granule dynamics in C9orf72-associated frontotemporal dementia and amyotrophic lateral sclerosis. Nature Medicine 24:1136. doi: 10.1038/s41591-018-0071-1

